# Effects of Concussion on the Relative Contributions of Sensorimotor Memories during Adaptation to Unpredictable Spring-Like Loads

**DOI:** 10.1101/2024.05.16.594491

**Authors:** Devon D Lantagne, Leigh A Mrotek, Sheikh I Ahamed, Scott A Beardsley, Andrew J Grove, David H Leigh, Carolyn S Smith, Danny G Thomas, Robert A Scheidt

## Abstract

We examined the extent to which concussion impacts how implicit sensorimotor memories are used to compensate for changes in hand-held loads during goal-directed reaching. Recently concussed individuals performed computerized cognition tests and a robotic test of sensorimotor adaptation on three occasions: as soon as possible after injury; after clearance to return to activity; three months after injury. Non-concussed individuals (controls) were tested at inter-session intervals mimicking concussed group intervals. During robotic testing, subjects grasped the handle of a horizontal planar robot while reaching repeatedly to a target. The robot exerted spring-like forces that changed unpredictably between trials; this allowed us to estimate contributions of implicit sensorimotor memories to trial-by-trial performance by fitting a computational model to the time series of reach errors and robot forces. Symptom severity varied considerably within the concussed group at the first session. Computerized cognition tests revealed longer reaction times in the concussed group relative to control group in Session 1 only. Concussed subjects likewise had slower reaction time in the reaching task during the first but not later sessions. Computational modeling found abnormally high values of effective limb compliance in concussed individuals relative to the control group in the first session only, but did not find group differences in how sensorimotor memories contribute to reach adaptation in any session. Analysis of control group models identified a practice effect affecting the memory coefficients that may have masked initial effects of concussion on how implicit memories contribute to sensorimotor adaptation. Although a practice effect and a heterogeneous concussed cohort preclude strong conclusions, our findings suggest procedural improvements that may decrease the robotic test’s sensitivity to practice effects and increase its sensitivity to concussion-related changes in how implicit memories contribute to sensorimotor adaptation to unpredictable hand-held loads during reaching.

## Introduction

Approximately 180,000 sport-related concussions (SRCs) are reported annually in youth contact sports [1]. This statistic is concerning because concussion can cause temporary loss or degradation of brain function [2]. The cornerstone of modern sport concussion management is the removal of the athlete from sports activity and returning only after the concussion has resolved [2–4]. Failure to remove from play and premature clearance are associated with serious short- and long-term complications of traumatic brain injury [5–7]. Clearance for return to play is commonly based on assessments of symptom resolution, which assumes honest, best effort responses to symptom questionnaires and to computerized testing tools such as the off-field Sport Concussion Assessment Tool (SCAT) and computerized tests such as the CogState Computerized Cognitive Assessment Tool (CCAT; CogState, Inc., Melbourne, Australia) or the Immediate Post-Concussion Assessment and Cognitive Testing (ImPACT; ImPACT Applications, Inc., San Diego, CA, United States). These computerized assessments are only clinically reliable when test results can be compared against baseline test results. Unfortunately, it is not uncommon for athletes to intentionally suppress performance during baseline testing (i.e., malingering) in order to hasten clearance for return to play should they be concussed later on. During baseline testing, individuals can increase reaction times, decrease response accuracy, or feign memory problems during explicit recall testing [8]. After a concussive incident, individuals can deliberately underreport symptoms [9,10] which can decrease the apparent severity of symptoms for in-person assessments post-concussion. Although computerized tests provide built-in integrity checks to detect malingering, between 30-50% of individuals may be able to suppress performance during baseline testing satisfying integrity checks [11–13]. An assessment technique that is more resistant to malingering could be of great practical utility in the evaluation of concussion severity and in monitoring the time course of its resolution.

The assessment of motor control has shown promise in assessing concussion. Concussion has been known to impact cognitive-motor integration [14] and sensorimotor adaptation [15]. What is unknown is the question of being resistant to malingering. In one recent study by Lantagne et al. [8], a robotic assessment of implicit memories used in sensorimotor adaptation of reach extent to unpredictable spring-like loads has demonstrated resistance to intentionally degraded kinematic performance. Under these conditions, motor adaptation predominantly employs an implicit process [16] since explicitly-recalled performance did not improve memory modeling accuracy compared to actual reach errors which were not recalled. If this process and memory resources are maintained implicitly, these resources should not be recallable or be altered consciously when subjects are instructed to intentionally suppress performance. In this study, subjects were asked to perform ballistic out-and-back reaches to a target while a robotic device resisted outward movement with spring-like loads that changed unpredictably between trials. Systems identification techniques were used to fit a limited memory fast-process sensorimotor adaptation model to the trial-by-trial series of performance errors and spring-like loads. Coefficients returned by the fit describe the relative contributions of sensorimotor memories to motor adaptation. In addition to performing the reaching task with their best effort, subjects were also instructed to suppress performance by emulating symptoms associated with concussion (i.e., moving slower, delayed reaction times, fatigue, etc.). Subjects were able to negatively impact reach performance as quantified by increased variability in reach errors, reaction times, and target capture times as well as moving slower to the target and waiting longer to respond to a movement cue. However, the relative contributions of implicit sensorimotor memories remained unchanged, suggesting these contributions of motor adaptation to unpredictable perturbations are maintained implicitly and are resistant to intentional intervention by the individual. Considering concussion has been known to impact working memory recall [2,17] and cognitive-motor integration [14,18,19], and that concussion assessment is vulnerable to malingering [11–13], we put forward an initial investigation of the effects of concussion on implicit memories of sensorimotor adaptation.

In the present study, we sought to determine the extent to which concussion impacts the relative contributions of sensorimotor memories used in motor adaptation of movement extent against unpredictable spring-like loads to the hand. To this end, we first determined if kinematic measures and the contributions of sensorimotor memories to motor adaptation were sensitive to concussion shortly after injury. We then determined if these measures resolved over time across three time points. The first visit occurred as soon as possible after injury; the second after subjects became asymptomatic; and the third 3 months after injury as a healthy baseline. CogState neurocognitive testing served as an objective measure of concussion and its recovery within the study cohort. The robotic assessment of implicit sensorimotor adaptation had subjects grasp the handle of a horizontal planar robot while performing ballistic out-and-back reaches to a target while the robot resisted outward movement with a spring-like force. Spring strength changed unpredictably between trials. The unpredictable perturbations to the hand elicited movement errors that engaged sensorimotor adaptation. Systems identification techniques were used to fit the trial-by-trial series of performance errors and spring strengths to quantify the extent to which sensorimotor memories contribute to motor performance. Concussion impacted kinematic outcome variables such as timing and accuracy measures. Most of these parameters were no different from controls by the second visit, except for time taken the capture the target which did not return to magnitudes of controls after three months. Concussion also impacted sensorimotor adaptation by changing how concurrent spring strength influenced subsequent reach errors. However, we did not find significant changes in how memories were used. Further conclusions within the first session were confounded by a practice effect that was present in the control group which masked any potential effects of concussion. The practice effect resolved by the second visit, indicating a requirement to perform a practice test.

## Methods

### Participants

Twenty healthy young adults [age: 21.8 ± 2.3 years (mean ± standard deviation, here and elsewhere); 11 females] and 22 recently concussed individuals (age: 20.1 ± 1.4 years; 14 females) provided written, informed consent to participate in this longitudinal study. Non-concussed individuals (i.e., control subjects) were recruited from the general Marquette community and none had any known neurological deficits or had been diagnosed with concussion within the previous 12 months. Concussed subjects were solicited from the Marquette University Medical Clinic. All subjects had intact proprioception and normal or corrected-to-normal vision. Institutional approval (HR-3233) was obtained for all experimental procedures in accord with the Declaration of Helsinki. Subject recruitment began February 1^st^, 2017, and ended June 1^st^, 2022.

All concussed individuals provided additional written consent allowing the study team to access medical records from the Marquette University Medical Clinic. These records included results from the Post-Concussion Symptom Scale (PCSS), which was obtained during the first clinical visit. The PCSS rates the initial severity of concussion by assessing 22 symptoms on a scale from 0 to 6, where the value 6 indicates highest severity. We used the total number of symptoms and the sum of all symptom severities as two composite measures of initial concussion severity.

### Experimental Setup and Procedures

We asked subjects to participate in three experimental sessions lasting approximately 45 minutes each.

Concussed subjects were asked to attend the first session as soon as possible after their injury, the second session after the clinic had cleared them to return to normal activity (typically when individuals were asymptomatic), and a third session three months after the injury. Control subjects were scheduled to mimic the inter-session intervals of the concussed group; they attended the second session about 10 days after the first and the third session approximately three months later.

Subjects performed two tests within each experimental session. The first was a clinically-accepted computerized assessment of cognitive performance (CogState, Inc, Melbourne, Australia). The second was a novel test of sensorimotor learning during goal-directed reaching [8,16,20].

#### Computerized Cognitive Assessment

Subjects were seated in a quiet room, where they completed four tasks from the CogState task library. All four tasks featured virtual playing cards that were displayed sequentially on a computer monitor. Subjects were to provide ‘yes’ or ‘no’ responses to a task-dependent question using the right and left computer mouse buttons, respectively. The first task, Detection (DET), assesses sensorimotor processing speed using a simple reaction time test: subjects were to press only the ’yes’ button as soon as possible after a card was displayed. The second task, Identification (IDN), assesses attention via choice reaction time: subjects were to indicate as soon as possible after each card presentation whether the card was red by pressing the appropriate mouse button. The third task, One-Back (ONB), additionally assesses declarative working memory: subjects were to press the appropriate mouse button to indicate whether the previous card was the same as the current card. The last task, One-Card Learning (OCL), also measures declarative working memory over a longer item span. Instead of recalling only the previous card, subjects were to press the appropriate mouse button to indicate whether the current card was previously shown at any time during the OCL task. The CogState software reports the mean button press accuracy and reaction time for each task.

#### Robotic Assessment of Sensorimotor Learning

Subjects sat in a high-backed chair and grasped the handle of a horizontal planar robot, which they moved with their right hand (Fig 1A). An opaque screen mounted just above the plane of movement occluded direct view of both their hand and the robot arm. A computer projected visual stimuli onto the screen; these included a home target, a goal target, and (occasionally) a small cursor that provided real-time visual feedback of hand position. The home position and the target position were indicated using 0.4 cm diameter white dots spaced 10 cm apart along the line intersecting the horizontal and sagittal planes passing through the shoulder center of rotation. The goal target was positioned farther from the subject on that line than the home target. When displayed, the hand’s cursor appeared as a small white 0.4 cm diameter dot that accurately tracked hand position.

**Fig 1.**
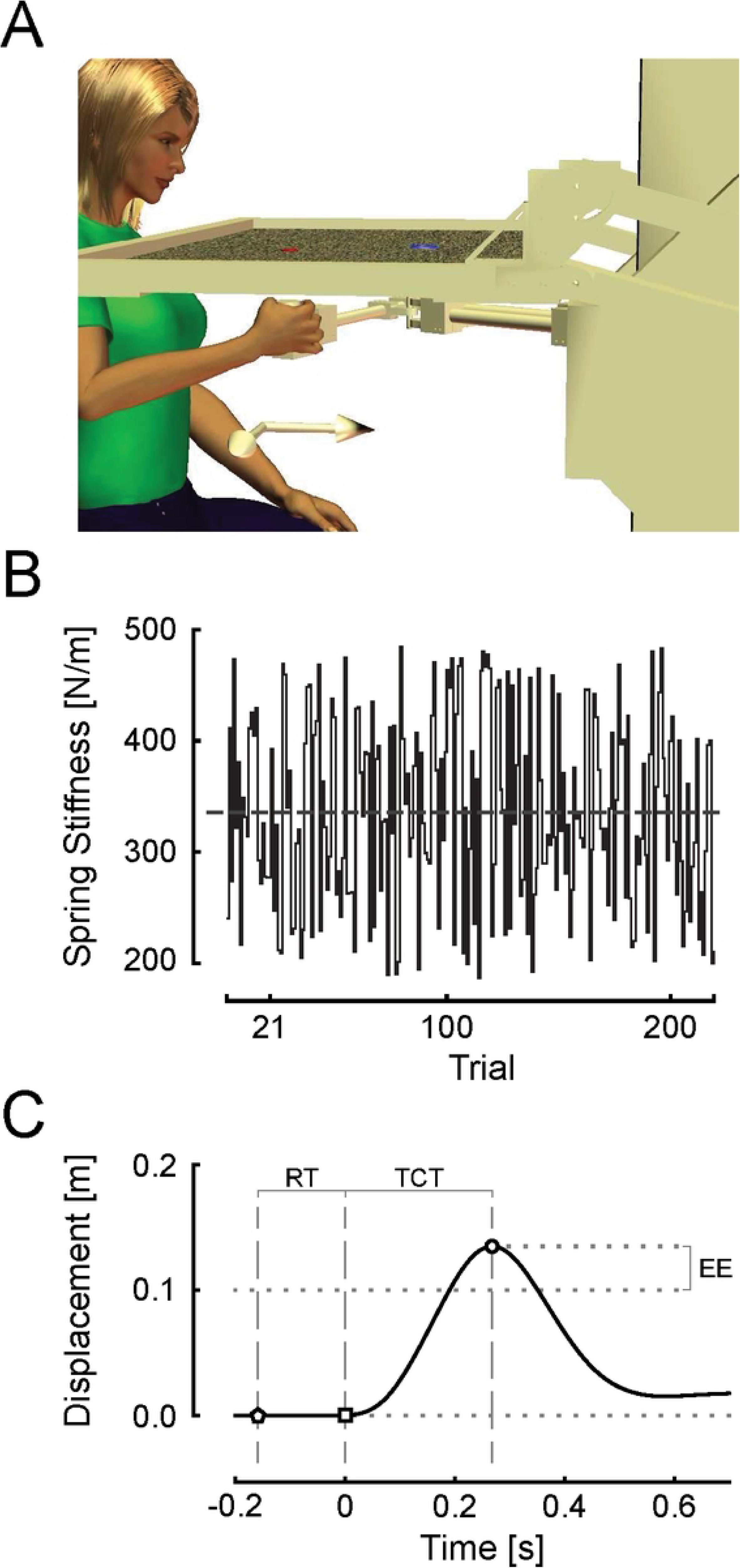
Setup and Measures. **A** Experimental setup: Home and goal targets were projected on a horizontal screen which occluded the subject’s view of the hand. **B** Trial-by-trial changes in spring stiffnesses rendered by the robot. Horizontal line indicates mean spring-like load of 335 N/m. **C** Hand displacement of a typical reach trial. Circle – point of maximum reach extent; square – point of movement onset defined as 10% of peak velocity; pentagon – point when the ‘GO’ cue was presented. RT – reaction time from GO cue to movement onset; TCT – target capture time from movement onset to peak extent; EE – reach extent error relative to the target.

We adapted a previously published approach [8,16] to quantify how implicit memories of performance contribute to sensorimotor adaptation of reaching movements. Briefly, subjects performed 220 trials of a goal-directed, out-and-back reaching task while holding the robot’s handle. During each reach, the robot applied a spring-like resistance that opposed movement away from the home target. This spring strength was changed pseudorandomly between trials (Fig 1B) such that subjects could not predict the next spring strength. The spring stiffnesses were drawn from a uniform distribution with a mean strength of 335 ± 86 N/m. The robot also applied a virtual force channel (c.f. [21]) to actively constrain all movements of the handle to the straight line between the home and goal targets. This aspect of the experimental design constrained kinematic performance errors to those of extent, not direction, thereby simplifying subsequent analysis (cf. [8,20]). The robot recorded hand position and applied hand forces at a rate of 1000 samples per second.

To begin a trial, subjects moved their hand to the home position. Cursor feedback was withheld until the hand was within 2 cm of the home target. After the hand was held there for 0.25 seconds, a GO cue signaled subjects to initiate a reach movement. All visual stimuli were removed at the time of the GO cue such that subjects had to reach to the remembered location of the target without cursor feedback. After the reach was performed, subjects were instructed to return to the home position to begin the next trial.

The first 20 reaching trials were considered “practice trials,” wherein the hand’s cursor was projected onto the screen throughout the entire movement to allow subjects to learn the spatial requirements of the reaching task. The remaining 200 trials were used as “test trials” and did not display concurrent visual feedback of hand motion. We used these trials to analyze kinematic performance and trial-by-trial adaptation of goal-directed reaching to a remembered spatial target.

### Data Analysis

#### Computerized Cognitive Assessment

We used the CogState Research2 toolkit to extract standardized measures of performance accuracy and reaction time for each of the four tests (DET, IDN, ONB, and OCL) performed in each session by each subject. The toolkit corrects for non-normality in the raw data by applying a log_10_-transform to response times and an arcsine-transform to the square root of the proportion of correct trials to total trials in each test [22–24]. All subjects’ test measures were standardized to the control group’s within-session performance using z-scores.

#### Robot assessment of sensorimotor learning

We used a semi-automated algorithm to verify that subjects successfully performed the task as instructed on each trial. We confirmed the algorithm’s recommendations through direct visualization of hand displacement, velocity, and acceleration profiles in each reaching trial. Trials were excluded from further analysis for any of the following reasons: if the hand drifted 1 cm or more from the home position before the GO cue; if the acceleration profile of the out-and-back reach was not triphasic (i.e., suggesting corrective movements or mid-reach pause); or if the reach was not directed along the channel constraint causing the robot’s motor torques to exceed safety limits (35 Nm).

We extracted three kinematic outcome variables from the measured hand displacement trace in each trial. *Reaction time* (Fig 1C: “RT”) was defined as the time between the GO cue (Fig 1C: pentagon) and movement onset (Fig 1C: square), defined as the moment when hand speed exceeded 10% of its first peak value. *Target capture time* was the difference between movement onset and the moment the hand reached the maximum extent of outward movement (Fig 1C: circle, “TCT”). We quantified movement accuracy using *reach extent error* (*ɛ*_*i*_), defined as the signed difference between maximum movement extent and the actual target distance of 10 cm (Fig 1C: “EE”). Across all test trials, we computed the mean and standard deviation for each outcome variable to yield six total measures of reaching performance.

#### Modeling of sensorimotor adaptation

We examined the extent to which recently sustained concussion alters the way individuals process sensorimotor memories of reach performance as they adapt to changing environmental loads. To do so, we used a simple computational model (Equation 1) that was previously found capable of quantifying the relative influence of prior reach extent errors (*ɛ*) and prior environmental spring strengths (*k*) on target capture performance during repetitive reaching [8,16,20,25]:

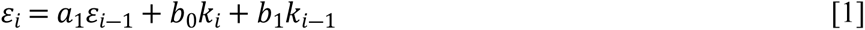

Prior to model fitting, the across-trial mean value for each of the factors is subtracted from their respective time series. In Equation 1, subscript *i* is the trial index of test trials ranging from 21 to 220. The term *ɛ*_*i*―1_is considered a proxy for implicit memory of the prior reach error because it is inaccessible to declarative recall even though it possesses significant power to predict performance on the following reach [16]. The impact of the robot’s physical resistance is represented by the current trial’s spring strength (*k*_*i*_) and the memory of the previous spring strength (*k*_*i*―1_). The best fit model coefficients, *a*_1_ and *b*_1_, describe the relative contributions of sensorimotor memories to reach performance on a trial-by-trial basis. The non-memory coefficient, *b*_0_, describes the physical influence of the spring on the hand (i.e., stiff springs result in smaller movements and lighter springs extended movements).

### Statistical Hypothesis Testing

We tested three primary hypotheses. We first hypothesized that the outcome measures described above would reflect measurable effects of concussion at the first testing session since it is most proximal to the date of the injury. To test this, we performed a-priori t-tests between groups within session 1. We next hypothesized that observed effects of concussion would resolve (i.e., become comparable to controls) by the second session. We tested this hypothesis using a-priori t-tests between groups within session 2. Finally, we hypothesized that recently concussed subjects would perform no different than controls by the third experimental visit (3 months after injury), reflecting resolution of all symptoms. Again, this hypothesis was tested using a-priori t-tests between groups within session 3.

After data collection, some outcome measures appeared to change systematically across sessions within the control group, indicative of a practice effect. To test for the presence of a practice effect (our secondary hypothesis), we performed one-way, repeated measures MANOVA across sessions within the control group to determine if and when outcome measures might systematically change.

All data processing and model fitting was performed using MATLAB 2022a (The MathWorks, Natick, Massachusetts). Statistical processing was performed using SPSS 28 (IBM corp. Armonk, New York). Statistical significance was set to a family-wise error rate of α = 0.05.

## Results

Seven of the 22 concussed subjects and two of the 20 control subjects did not attend the third experimental session three months out from initial injury. Because we intended to consider measured performance in the third session as a healthy baseline, these subjects were excluded from further analysis. One additional concussed subject was excluded from further consideration because they sustained a second concussion between the first two sessions. In total, 18 healthy subjects and 14 recently concussed subjects were considered for further analysis. No concussed subjects reported worsening symptoms during or after performing the experiments.

### Clinical Assessment of Post-Concussion Symptoms

The concussed group was heterogeneous with respect to the duration between initial injury and the first experimental session (Fig 2A), the number of symptoms as indicated on the post-concussion symptom scale (PCSS) (Fig 2B), and the sum of PCSS responses (Fig 2C). The time between injury and the first session ranged from 0 to 17 days, (average: 5.9 ± 4.1 days). The number of symptoms reported on the PCSS ranged from 9 to 22 symptoms across the study cohort (average: 17.8 ± 4.2 symptoms). The severity of these symptoms also varied markedly across the cohort; the sum of PCSS symptom scores ranged from 16 to 105 on a scale that has a maximum symptom severity score of 132 (cohort average: 59.0 ± 26.2).

**Fig 2.**
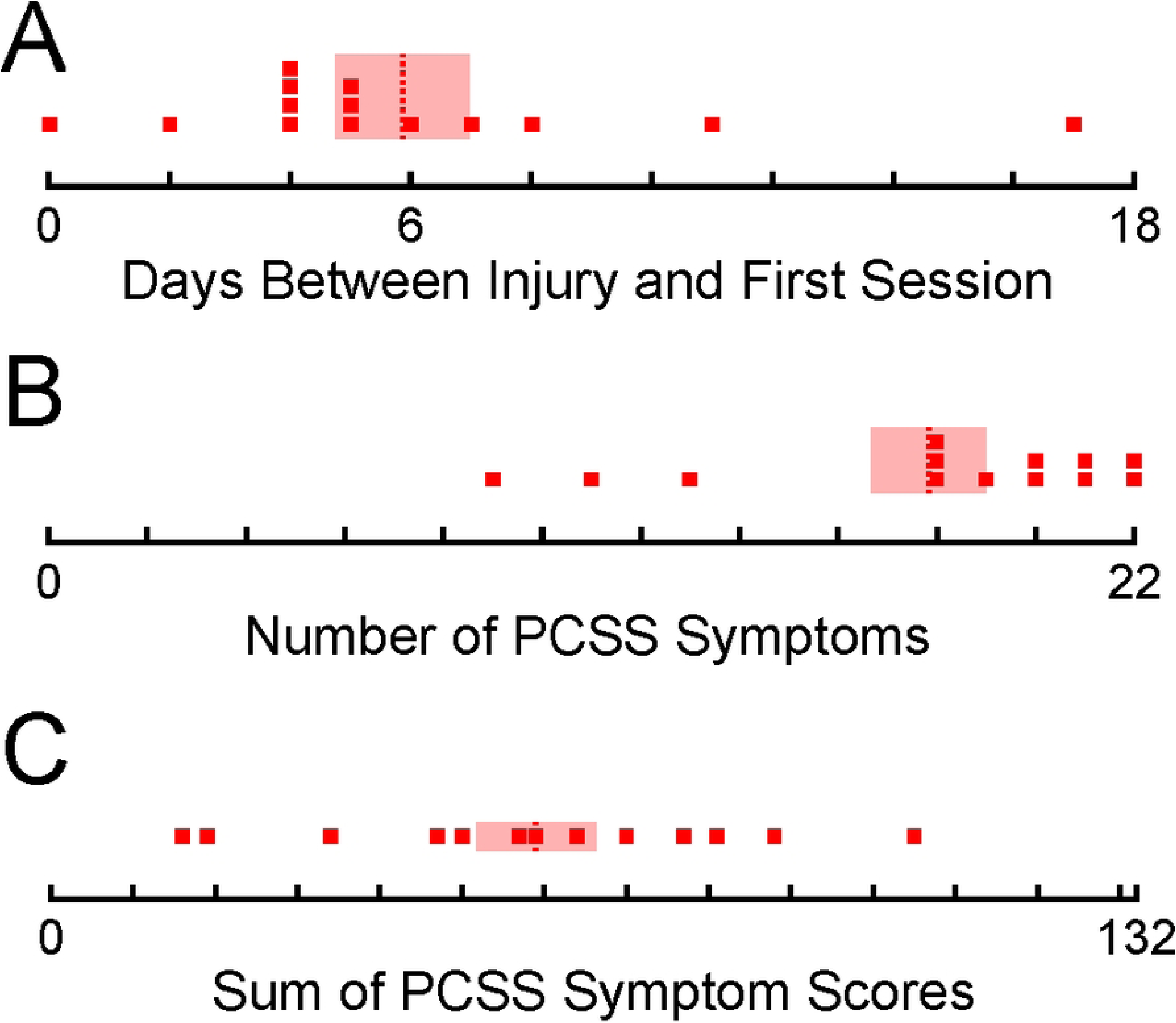
Concussed cohort data. **A** Delay between injury and attending first session. Red dashed line: group mean; transparent red rectangle: ± 1 SEM. Post-concussion symptom scale (PCSS) **B** number of reported symptoms (out of 22) and **C** symptom score grand total (out of 132).

### Computerized Testing of Cognitive Performance – Session 1

We next assessed cognitive performance as quantified by response accuracy (Fig 3A) and reaction time (Fig 3B) for each CogState task at the time of the first experimental session. Accuracy was assessed differently for each task: DET) a response was deemed accurate only if it occurred after a card was presented; IDN) a response was deemed accurate only if it correctly identified whether a card was red or black; ONB) a response was deemed accurate only if it correctly indicated whether the presented card was the same as the last card; and OCL) a response was deemed accurate only if it correctly indicated whether the presented card had been previously presented during the OCL task. In all cases, reaction time was quantified as the amount of time between the presentation of a card and the subject’s response. In Figure 3, all performance measures are presented in z-scores (i.e., standardized to the performance of the control group).

**Fig 3.**
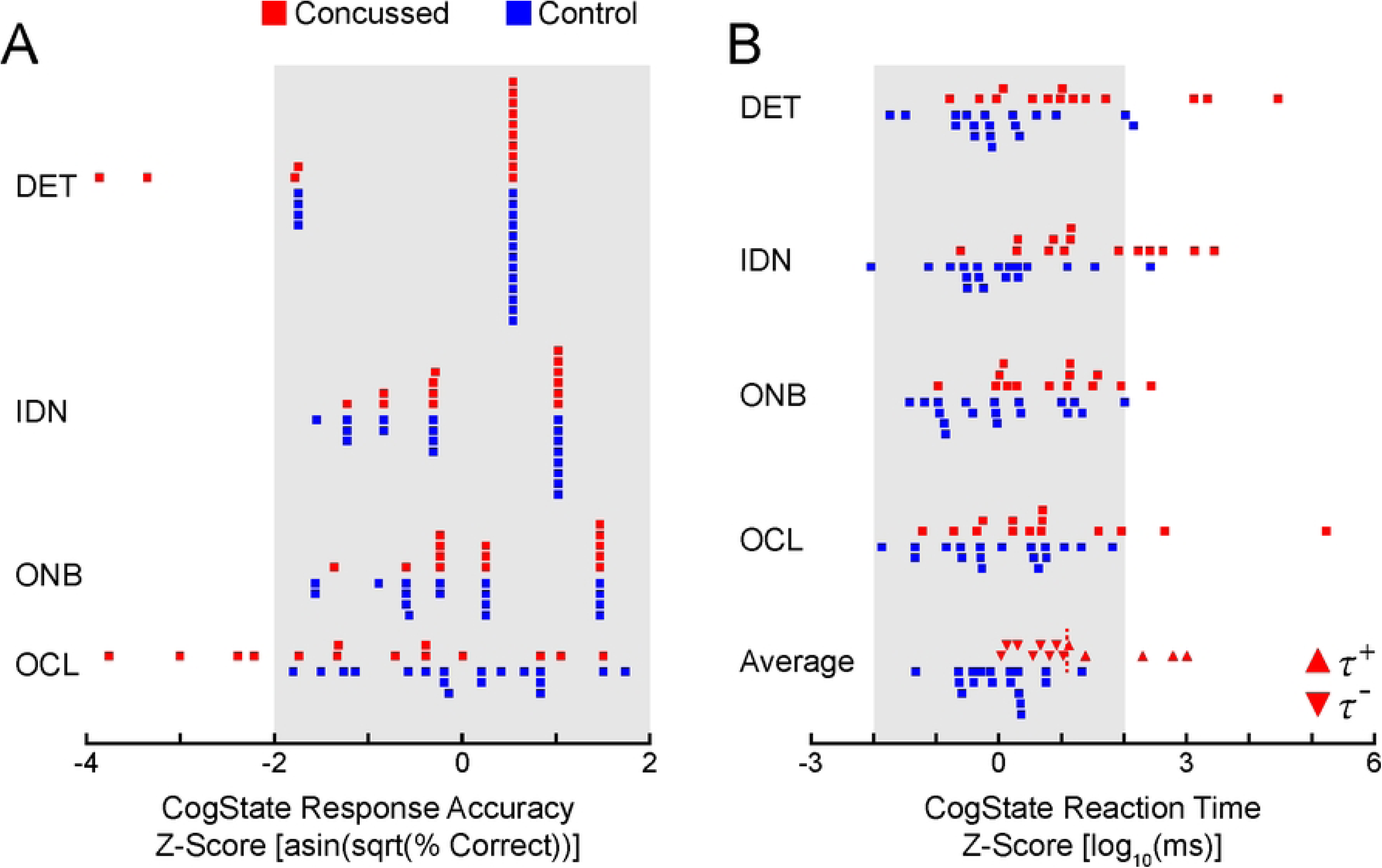
Z-scores of transformed CogState response accuracies. **A** and reaction times **B** for each task. DET: Detection; IDN: Identification; ONB: One-Back Memory Recall; OCL: One Card Learning; and Average: average score across tasks. Concussed group split between the ***Ꚍ***^+^(above-average) and ***Ꚍ***^―^(below-average) average reaction time score. Vertical dotted red line: across-subject average reaction time of concussed group. Grey box: 2 standard deviations above and below the mean (95% CI). Positive accuracy scores and negative reaction time scores indicate better performance relative to controls.

With respect to response accuracy in session 1 (Fig 3A), only the OCL task showed differences between groups, with concussed subjects responding less accurately than controls on average. The other three tasks (DET, IDN, and ONB) did not exhibit group differences. In these three tasks, scores from the concussed subjects rarely deviated more than two standard deviations from the mean of the control group (Fig 3A; compare the number of red squares within/without the gray shaded regions). By contrast, reaction times appeared to be more sensitive to concussion (Fig 3B); the DET, IDN, and ONB tasks all showed significant differences between groups (all T_30_ > 2.309, p < 0.028). The groupwise difference in the OCL task did not quite achieve significance (T_30_ = 1.820, p = 0.079). With respect to reaction times, a few more concussed subjects yielded scores outside the range spanned by the control group’s mean ± 2 standard deviations; nevertheless, most concussed subjects’ scores remained within the “normal” range of controls.

Given that reaction times appeared more sensitive to concussion than accuracy scores at the time of session 1, we computed the average response time z-score across tests within each subject and then compared these averaged scores between groups (Fig 3B: *Average*). Again, we noted substantial overlap in performance between the concussed subjects and the controls. On average, all but three concussed subjects performed within 2 standard deviations of control mean. Despite this overlap, both groups were found to be significantly different from one another (T_30_ = 3.882, p < 0.001).

Considering the overlap of timing and accuracy scores and the extended duration between the injury and the first visit, we suspected that some concussed subjects may have significantly recovered before the first session which could contaminate any effect of concussion in subsequent analyses. We subdivided the concussed group into two concussion severity groups based on each subject’s CogState averaged reaction time score relative to the concussed cohort mean. One subgroup, *τ*^―^, was defined as being below the mean processing speed (N = 9; Fig 3B red downward triangles) while a second subgroup, *τ*^+^, was defined above the mean (N = 5; Fig 3B red upward triangles). The PCSS symptom sum and symptom count could also be considered as a method to divide the concussed group based on initial severity. However, correlation analyses between the PCSS against CogState timing and accuracy measures revealed that the participant’s self-reported PCSS did not have any predictive power on the more objective CogState measures (all p > 0.205). We therefore discontinued using the PCSS for the remainder of this study.

### Comparison of Concussion Subgroups

We explored the impact of creating two concussed subgroups based on average cognitive processing speeds. Specifically, we wanted to determine if reaction time or accuracy scores across the four CogState tasks were different between groups. To do this, we performed a one-way MANOVA for reaction time and accuracy scores, separately, within session 1 between the controls and the two concussed subgroups. The analysis of response accuracy did not reveal a significant effect between groups [F_(8,52)_ = 2.045, p = 0.059], although this is marginal. However, analysis of reaction time did reveal a significant difference between groups [F_(8,52)_ = 4.748, p < 0.001]. This result warranted subsequent univariate tests within each CogState task to identify which tasks were sensitive to concussion, and if so, which group was different. P-values of univariate tests were Bonferroni corrected for four multiple comparisons. Subject group had a significant effect on reaction time for the DET and IDN tasks (assessing simple and choice reaction times) [DET: F_(2,29)_ = 14.720, p < 0.001; IDN: F_(2,29)_ = 16.108, p < 0.001]. In all significant cases, post-hoc Dunnet t-tests indicated only concussed subjects with long reaction times (*τ*^+^ group) were different than controls (all p < 0.001) whereas the concussed subjects with shorter reaction times (*τ*^―^ group) were not significantly different from controls in any task (all p > 0.062). The other two tasks, ONB and OCL which assess declarative working memory and learning, did not yield significant effects of group after multiple comparisons [ONB: F_(2,29)_ = 3.627, p = 0.157; OCL: F_(2,29)_ = 4.581, p = 0.075].

These results highlight two findings: 1) that the reaction time measure from the Detection and Identification tasks were sensitive in detecting cognitive deficits in our subject sample at the time of the first experimental visit whereas the One-Back memory and One-Card Learning tasks did not show differences; and 2) that concussed subjects with shorter average reaction times did not perform differently than controls, whereas concussed subjects with longer average reaction times demonstrated clear reaction time deficits. To mitigate the concern of including potentially recovered individuals as members of the concussed cohort, we excluded the *τ*^―^ group from further analyses because their performance was not different than that of healthy controls.

### Cognitive Function Across Time

With the newly organized *τ*^+^concussion cohort, we investigated our three primary hypotheses: 1) that concussed subjects will exhibit measurable differences in outcome measures compared to controls in the first session; 2) that those differences may resolve by session two; and 3) that any remaining differences will resolve by session three. Each of these three hypotheses were tested using a-priori t-tests on each CogState task’s reaction time z-scores.

Concussion resulted in reaction time deficits as quantified by the Detection task’s measure of simple reaction time (Fig 4A). During the first visit, concussed subjects took significantly longer to react than controls (T_21_ = 4.993, p < 0.001). By the second visit, recently concussed subjects still took longer to react than controls despite being cleared to return to activity (T_21_ = 2.595, p = 0.017). However, by the third visit three months after the injury, the concussed group did not perform differently than controls (T_21_ = 1.114, p = 0.278).

**Fig 4.**
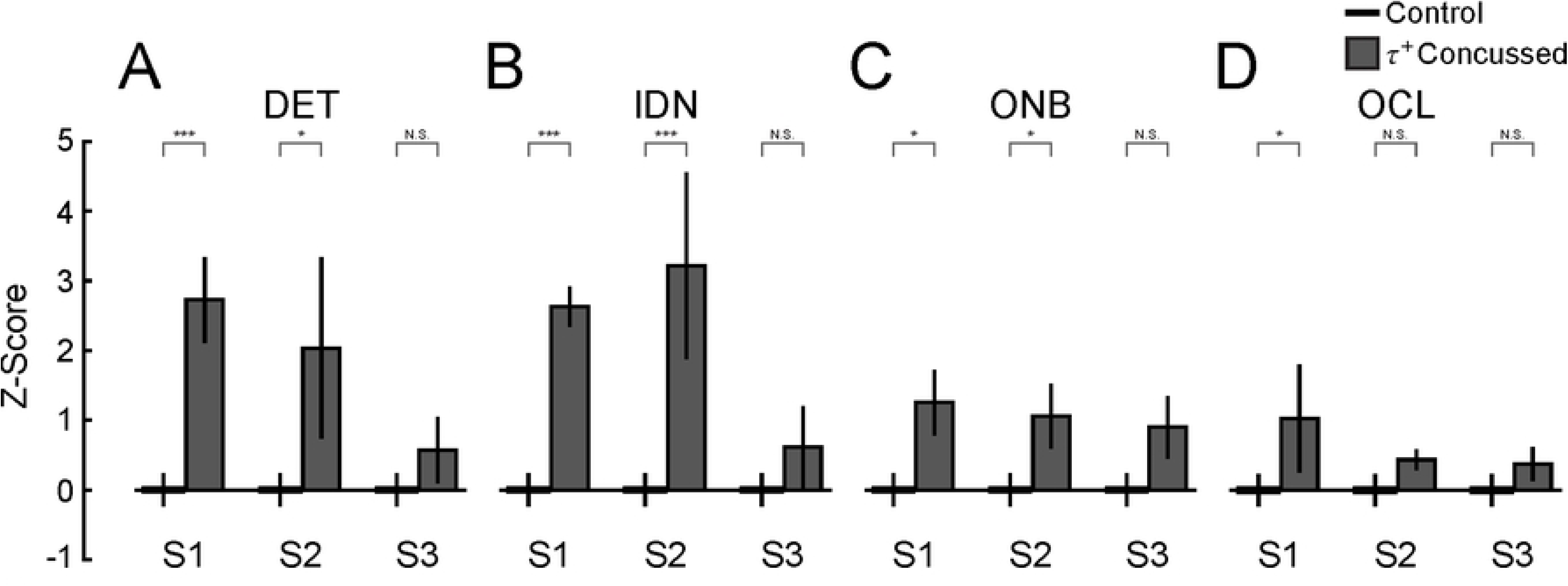
Comparison of CogState z-scores between control and ***Ꚍ***^+^ concussed cohorts across sessions. Z-scores computed relative to control group within session. Control cohort is standardized to itself (and thus the mean of their z-scores equals zero). Error bars: ± 1 SEM. Significance bars: a-priori t-tests for primary hypotheses. Significance bars: N.S. not significant; * p < 0.05; ** p < 0.01; *** p < 0.001.

The Identification task identified deficits in choice reaction time (Fig 4B). Concussed subjects took longer to respond during the first session (T_21_ = 5.518, p < 0.001). By the second visit, concussed subjects did not perform differently from controls after adjusting for unequal variances between groups (T_4.3_ = 2.371, p = 0.073). By the third visit, both groups performed similarly (T_21_ = 1.134, p = 0.270).

Concussion also resulted in timing deficits of the One-Back working memory task (Fig 4C). Concussed subjects took longer to respond during the first visit (T_21_ = 2.454, p = 0.011). At the time of the second visit, concussed subjects still performed worse than controls, albeit marginally (T_21_ = 2.086, p = 0.049). Three months after injury, concussed subjects performed comparable to controls (T_21_ = 1.780, p = 0.090).

Lastly, unlike the preceding three tasks, the One-Card learning task’s reaction time was not significantly impacted by concussion relative to controls in our cohort (Fig 4D). Concussed subjects did not perform significantly different from controls during the first session after adjusting for unequal variances between groups (T_4.4_ = 1.764, p = 0.146). Performance between groups was also comparable during the second visit (T_19.3_ = 1.556, p = 0.136) and third visit (T_21_ = 0.809, p = 0.428).

These results highlight three main findings: 1) Concussed subjects in the *τ*^+^group possessed reaction time deficits at the time of the first session in tasks of simple reaction time (DET), choice reaction time (IDN), and one-back declarative working memory (ONB); 2) the IDN and ONB tasks reported differences between groups at the second session when concussed subjects typically present asymptomatically, demonstrating that some cognitive deficits resolve slower than symptoms; 3) all four cognitive tasks report no differences between groups at the third visit, suggesting the concussed cohort may have recovered within three months after their injury. With this clinically-approved assessment of concussion reporting deficits in our concussed cohort, we next investigate the effects of concussion on reaching kinematics during the robotic assessment of sensorimotor adaptation.

### Robotic Reach Testing: Kinematic Performance

Kinematic data from selected control and concussed subjects are shown in Figures 5A and 5B. Reaches of the control subject (Fig 5A) were consistent with ballistic out-and-back movements (target capture time: 201 ± 8 ms). This subject consistently overshot the target (reach error: 3.2 ± 1.8 cm) which is typical when visual feedback is not presented [16,20]. A concussed subject is shown in Figure 5B. These movements were also ballistic, but with longer and more variable target capture times (326 ± 29 ms) and greater reach errors (7.2 ± 1.8 cm). Figures 5C and 5D show each subject’s reach errors can be explained in part by a contribution from the spring strength felt by the hand. Aside from the vertical bias representing the mean reach error (which we subtract out for modeling analysis), the concussed subject’s reach error as a consequence of the spring-like load applied to the hand (i.e., slope of the regression) was greater than the control subject (control: -0.88 cm^2^/N; concussed: -1.47 cm^2^/N).

**Fig 5.**
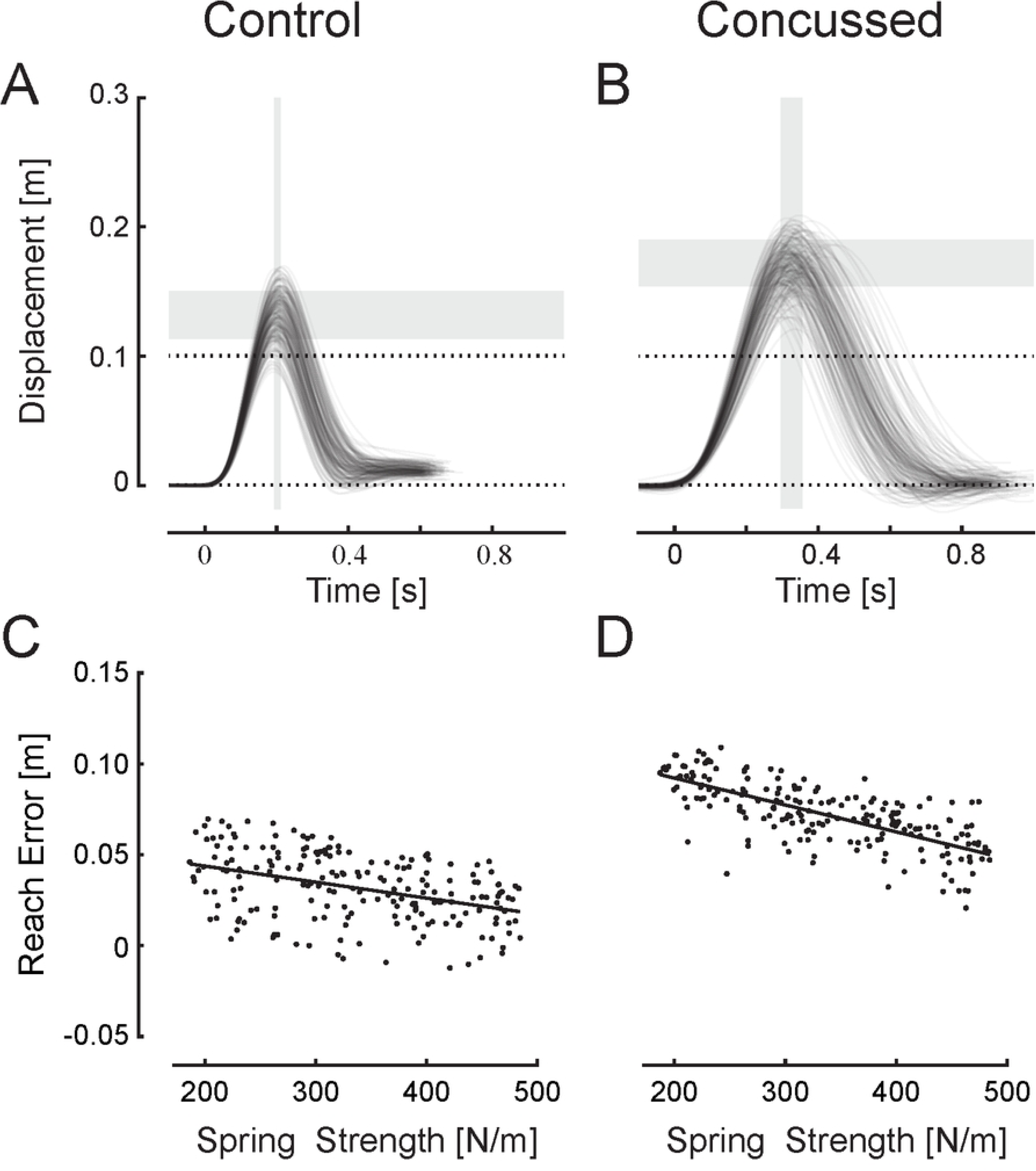
Kinematic performance. Reach trajectories for a selected control participant **A** and a selected concussed participant **B**. All traces aligned to movement onset (t = 0 seconds). Horizontal dashed line at 10 cm: target location. Horizontal grey bars: ± 1 SD about the mean reach error. Vertical grey bars: ± 1 SD about the mean target capture time. Reach error on each trial as a function of robotic spring stiffness for the same control participant **C** and the same concussed participant **D**. Solid black lines: best fit linear regressions.

We tested the three primary hypotheses for reach kinematic outcome measures (Fig 6). Kinematic timing measures were not normally distributed, so we performed a log_10_-transform on the data to satisfy the assumption of normality. However, three subjects (one concussed and two controls) yielded mean reaction times greater than 1000 ms or variability greater than 400 ms. These subjects were significant outliers and were removed from analysis of only reaction time measures. In the first session, subjects with more significant initial CogState performance deficits (*τ*^+^group) also had longer mean kinematic reaction times than controls (T_18_ = 2.220, p = 0.039). These differences resolved by the second session (T_18_ = 0.893, p = 0.384) and remained similar by the third session (T_18_ = 0.845, p = 0.409). Similarly, the variability of reaction times across trials was marginally greater (less consistent) than controls during the first session (T_18_ = 2.125, p = 0.048) which resolved by session two (T_18_ = 1.178, p = 0.254) and remained similar to controls during session three (T_18_ = 1.061, p = 0.303). Together, concussed individuals with extended reaction times during computerized cognitive tasks resulted in longer and more variable kinematic reaction times when tested shortly after injury, but these deficits resolved by the time subjects reported minimal symptoms and were cleared to return to activity. Repeating the analysis when including the three outlying subjects produced a similar pattern of results.

**Fig 6.**
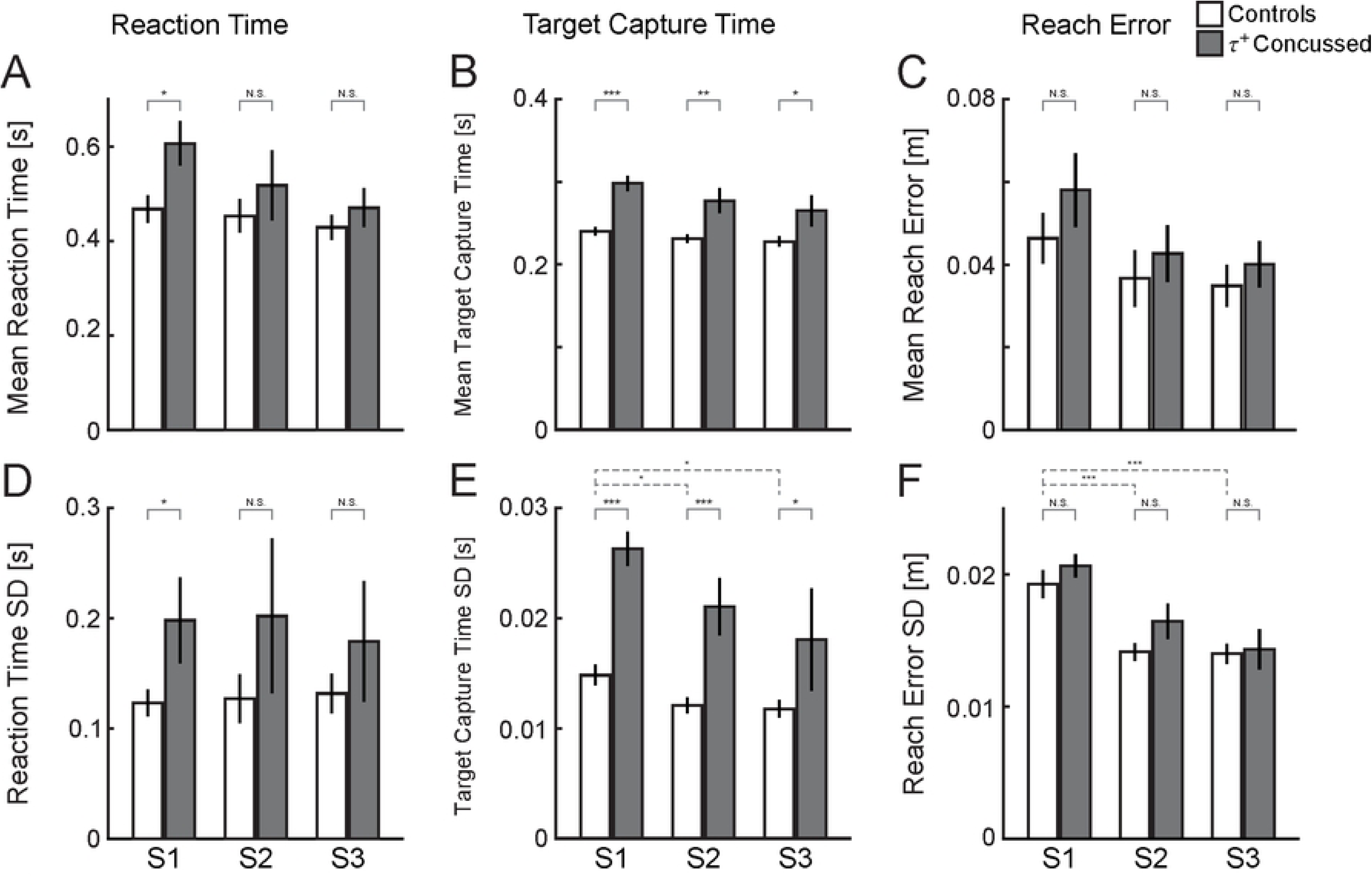
Comparison of kinematic measures of reaching performance between control and ***Ꚍ***^+^ concussed groups. *Column 1*: reaction time. *Column 2*: target capture time. *Column 3*: reach extent error. *Top row*: mean performance across sessions. *Bottom row*: precision of performance across sessions. Error bars: ± 1 SEM. Solid significance bars: primary hypotheses via a-priori t-tests between groups. Dashed significance bars: secondary hypothesis via post-hoc comparisons of a-posteriori repeated measures, one-way ANOVA within the control group. N.S. not significant; * p < 0.05; ** p < 0.01; *** p < 0.001.

We next tested the three primary hypotheses for target capture time – the duration between movement onset and the maximum extent of the reach (Fig 6 B and E). These measures were also transformed using a log_10_-transform. The concussed group took longer to capture the target during the first visit (T_21_ = 4.784, p < 0.001) and continued to show differences from controls during the second visit (T_21_ = 3.507, p = 0.002). Surprisingly, the concussed group continued to show deficits by the third visit three months after the initial injury (T_21_ = 2.357, p = 0.028). This pattern of results was also observed for the variability of target capture times between groups (S1: T_20_ = 4.245, p < 0.001; S2: T_20_ = 4.198, p < 0.001; S3: T_20_ = 2.107; p = 0.048). Most notable is that the target capture time of concussed subjects did not resolve toward values of controls even after three months from the injury.

Lastly, we tested the primary hypotheses on reach error performance (Fig 6 C and F). Unlike timing measures, reach errors were not impacted by concussion during any session as quantified by the mean or variability of reach errors (across all six comparisons: T_21_ < 1.065, p > 0.123).

The results of kinematic reaching measures highlight three findings: 1) that timing measures of reaction time and target capture time were sensitive to concussion at the first visit; 2) while the reaction time of concussed subjects became comparable to controls by the second session, target capture time did not return to performance like that of controls even by three months after the injury; and 3) that reach error performance did not show significant differences between concussed and control subjects at any time. However, the lack of detectable differences in reach errors across all trials does not capture trial-by-trial adaptive behavior; this will be investigated next by modeling reach performance using sensorimotor memories and comparing these relative contributions of motor memories within each of the experimental sessions.

### Robotic Reach Testing: Contributions of Sensorimotor Memories

Each subject’s series of reach errors and experienced spring strengths were fit by the sensorimotor adaptation model of Equation 1 which returned the relative contributions of sensorimotor information used to guide subsequent reaches (Fig 7). We first tested the three primary hypotheses on the relative contribution of the prior reach error (coefficient *a*_1_; Fig 7A) which represents a contribution from a sensorimotor memory. During the first experimental visit, the contribution of the memory of reach error was not different between the concussed and control groups (T_21_ = 1.113, p = 0.278) and remained similar between groups for the second session (T_21_ = 0.532, p = 0.600). During the third session, one concussed subject yielded an *a*_1_ coefficient significantly lower than any subject across all sessions (Fig 7A: X). Including this subject drives an inappropriate effect between the concussed and control groups within session three and was thus removed for this comparison. After removing this outlying subject, the concussed and control groups during the third session were not significantly different between groups (T_20_ = 1.866, p = 0.077). Ultimately, there was not a significant effect of concussion on the relative contribution of the prior reach error in our study cohort of concussed individuals.

**Fig 7.**
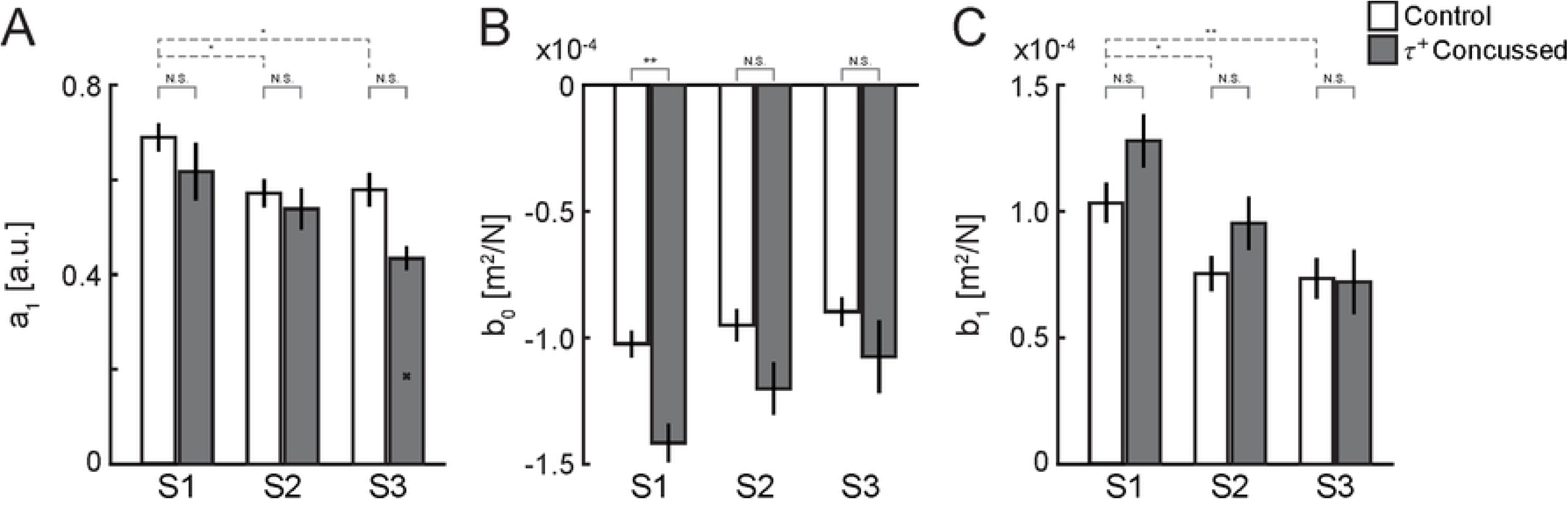
Comparison of sensorimotor adaptation model coefficients (Eqn 1) across sessions. **A** ***a***_**1**_: relative contribution of the previous reach error. Session three ‘x’: omitted outlying subject. **B** ***b***_**0**_: relative contribution of current spring stiffness. **C** ***b***_**1**_: relative contribution of previous spring stiffness. Error bars: ± 1 SEM. Solid significance bars: primary hypotheses via a-priori t-tests between groups. Dashed significance bars: secondary hypothesis via post-hoc comparisons of a-posteriori repeated measures, one-way ANOVA within the control group. N.S. not significant; * p < 0.05; ** p < 0.01; *** p < 0.001.

We next investigated the effects of concussion on the contribution from the series of spring strengths.

Concussion significantly increased the contribution of the concurrent spring strength (coefficient *b*_0_; Fig 7B) on reach performance during the first visit relative to controls (T_21_ = 3.655, p = 0.001) which mostly resolved by the second session (T_21_ = 1.897, p = 0.072) and was quite comparable to controls by the third session (T_21_ = 1.359, p = 0.189). We then tested the effects of concussion on the memory of the prior spring strength (coefficient *b*_1_; Fig 7C) within each of the three sessions but, similar to the memory of the prior reach error, no differences were observed between the concussed and control groups in any session (all sessions T_21_ < 1.524, p > 0.143).

### Practice Effects

While investigating kinematic performance, we noticed the control cohort displaying different values of outcome measures across sessions (e.g., Fig 6F: controls) and could be considered a practice effect through repeated participation in the robotic assessment of sensorimotor adaptation. While this potential effect does not diminish prior findings when comparing groups within session, the presence of a learning effect will be important to characterize if the robotic assessment of sensorimotor adaptation is to be used in a clinical setting.

We therefore formed a secondary hypothesis to determine if outcome measures within the control group varied across sessions. To test this, we performed a one-way, repeated measures MANOVA across sessions for kinematic outcome measures and a separate MANOVA for modeling coefficients.

We first determined if any of the six kinematic outcome variables contained session effects within the control group. A MANOVA identified a significant effect in at least one of the outcome variables [F_(12,50)_ = 3.917, p < 0.001]. Subsequent univariate tests for each of the six kinematic outcome variables, after adjusting for six comparisons, revealed the variability of target capture time (Fig 6E) and the variability of reach errors (Fig 6F) possessed a significant session effect [both F_(2,30)_ > 10.940, p < 0.002]. For both measures, post-hoc pairwise comparisons revealed that the first session was different from subsequent sessions (all p < 0.002). These results indicate a practice effect when measuring the variability of target capture time and reach error – subjects perform differently on the first session but reach consistent performance by session 2 onwards.

We then repeated the MANOVA for the three sensorimotor modeling coefficients which identified a significant effect of session in at least one of the coefficients [F_(6,64)_ = 2.921, p = 0.014]. Univariate tests identified significant effects of session on the contribution of the prior reach error (*a*_1_, Fig 7A) [F_(2,34)_ = 5.082, p = 0.035] and the prior spring strength (*b*_1_, Fig 7C) [F_(2,34)_ = 8.656, p = 0.003] but not the concurrent spring strength (*b*_0_, Fig 7B) [F_(2,34)_ = 3.238, p = 0.156]. In both cases, post-hoc comparisons revealed the first session was different than sessions two and three (all p < 0.029). These results suggest that both the memory terms in the sensorimotor adaptation model appear to exhibit a practice effect that reaches a steady state by the second session.

To provide a visualization of adaptive behavior from this practice effect, fitted coefficients of Equation 1 can be used to simulate a response from a select series of spring strengths. On such simulated input is a step increase (Fig 8: top left) where an absence of spring strength is followed by several trials of constant spring strength. Another simulated input is an impulse (Fig 8: top right) where sequential “on” and “off” perturbations occur. The model’s response to the step input (Fig 8: bottom left) for the first session yielded the greatest transient error concurrent with the initial perturbation exposure (trial 0) but reached steady state with minimal error. By contrast, the responses of subsequent sessions yielded smaller transient errors but greater steady-state errors. These characteristics are also present for an impulse input (Fig 8: bottom right). These simulations suggest that subjects adjusted how memories were used so as to reduce reach sensitivity to transient perturbations at the cost of increased steady-state error as training progressed.

**Fig 8.**
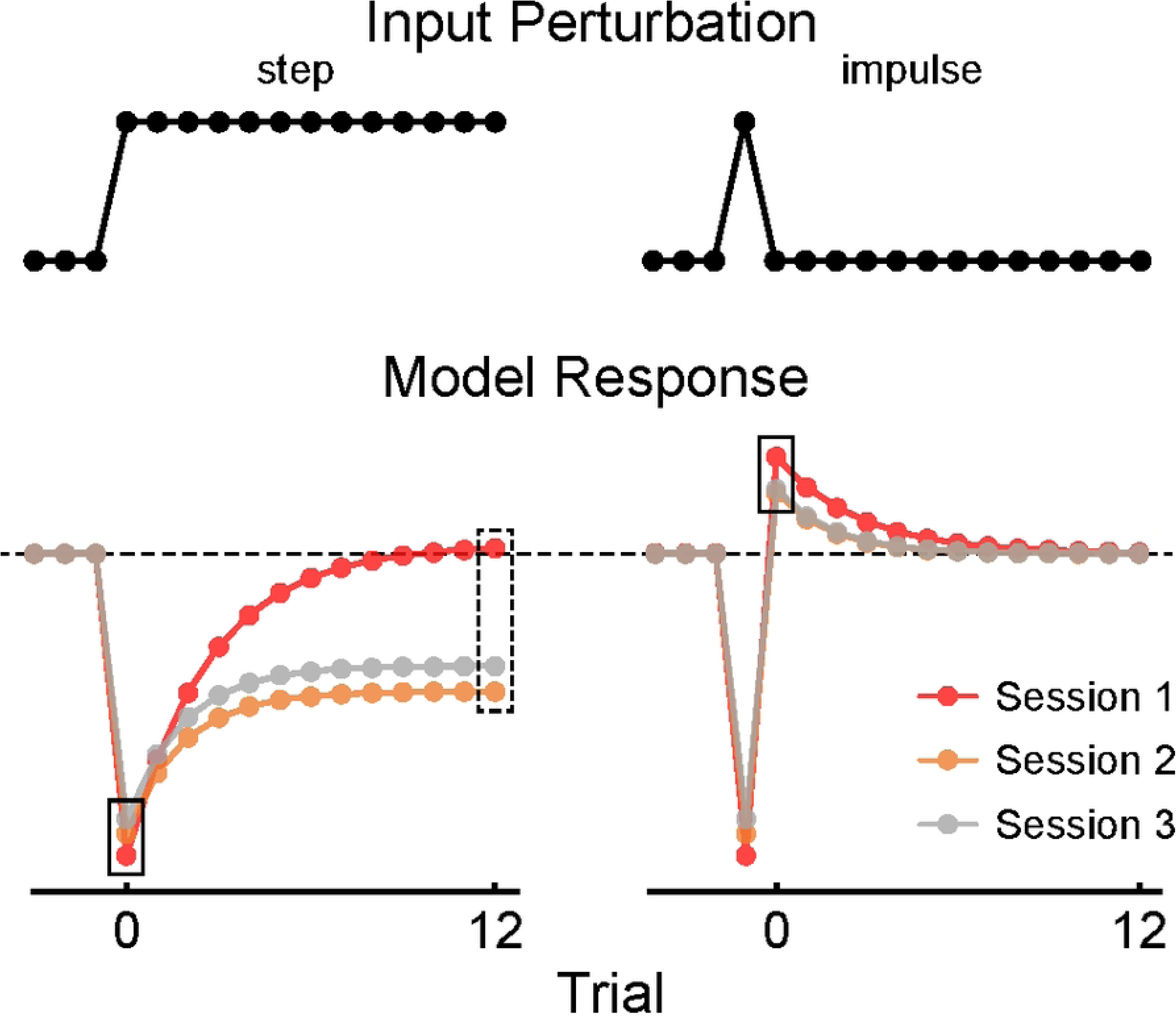
Memory model response analysis to step and impulse inputs (Eqn 1). *Top*: input series of spring stiffness (***k***) for a step and impulse increase. *Bottom*: model response (***ε***) for each session of the control group. Black rectangles: initial transient errors concurrent with the last change in perturbation. Dashed black rectangle: steady-state errors. Note the response of Session 1 yielded greater accuracy (smaller steady-state error in response to the step perturbation) at the expense of reduced precision (greater transient error in response to the impulse perturbation), whereas Sessions 2 and 3 yielded greater precision in response to the impulse perturbation but decreased accuracy in response to the step perturbation.

## Discussion

We determined the extent to which concussion impacts the relative contributions of sensorimotor memories to a goal-directed reaching task that required motor adaptation of movement extent to unpredictable spring-like loads applied to the hand. We recruited recently concussed participants to participate in three experimental sessions on separate days: once as soon as possible after injury, once when clinically cleared to return to activity, and once for a 3-month follow-up. Non-concussed control participants attended visits timed to match the average interval between concussed participant testing sessions. During each visit, each subject performed computerized cognition testing and a robotic test of goal-directed reaching. Notably, our sample of concussed subjects was a heterogeneous group. The concussed cohort attended the first experimental visit with variable latency after injury and with a broad range of symptom severities. Computerized cognition testing also revealed substantial overlap between the concussed and control groups in terms of cognitive reaction time. Consequently, we divided the concussed group into two groups: individuals with above- and below-average reaction time scores across our set of four cognitive tests. Concussed participants with below-average reaction times did not perform differently than the control group and were therefore removed from further analyses. Those with above-average reaction times in the first session of cognitive testing also had longer reaction times in the second session at which time symptoms should have been resolved. However, reaction time of the concussed group was comparable to controls by the third visit. Analysis of kinematic performance during robotic reach testing were also sensitive to concussion during the first visit. Notably, the time taken to capture the target was longer for the concussed group across all three time points, indicating motor performance deficits from concussion can exist several months after the injury. Sensorimotor memory modeling of reach error performance identified the contribution of the concurrent spring strength was sensitive to concussion during the first visit and was comparable to controls by the second visit, indicating concussion impacted how the brain adapted to unpredictable perturbations. However, the contributions from motor memories were not significantly different between the concussed group and controls in our small study cohort. Additionally, a practice effect was identified in the outcome measures of reach kinematics and in the contributions of sensorimotor memories. This practice effect was only observed between the first and second sessions, indicating longitudinal performance had achieved steady-state by the second visit and onward. Simulating step and impulse inputs to the memory model for each session revealed subjects adapted differently between the first and second session; with the first session’s response yielding larger transient errors and smaller steady-state errors compared to subsequent sessions. Future studies may wish to employ an extra testing session to mitigate practice effects and disentangle them from the effect of concussive brain injury on sensorimotor adaptation. Increasing task difficulty or changing sensory information (i.e., providing visual feedback or changing the statistics of the perturbations) may also impact how memories are used in sensorimotor adaptation – which may or may not improve effect sizes between recently concussed and healthy controls.

### Effects of Concussion on Sensorimotor Control and Adaptation

Concussion has been shown to result in motor control deficits that can manifest as longer reaction times [19,22,26], deficits in working memory and recall [17,22], increased variability of motor performance [18,27,28], and a decrease in motor adaptation learning rates [15]. In recent efforts to detect the presence of concussion history in athletes, Sergio and colleagues have developed an assessment of cognitive-motor integration using a portable tablet that captures the kinematics of finger pointing to a target (see [14] for review). The authors devised a reaching task that employed a 180-degree visuomotor rotation within the vertical plane of movement to dissociate the movement between the finger and a visual cursor [27]. This rotation requires individuals to employ an explicit cognitive rule-based strategy to counter the rotation (i.e., “if I need to move the cursor up, I must reach down to counter the 180-degree rotation”). This explicit cognitive transformation (a ‘level of dissociation’) exacerbates sensory integration deficits from concussion [27]. To further increase cognitive load and subsequently improve the detectable effect of concussion, a second tablet device oriented in the horizontal plane was used to record finger movement while the vertical tablet displayed rotated visual feedback, thereby introducing another spatial transformation to act as a second level of dissociation. The brain adapts to the second dissociation via implicit spatial recalibration (as when learning to use a computer mouse after using a touch tablet). Performing with two levels of dissociation yields larger effect sizes between previously-concussed and non-concussed individuals when measuring movement time and reach error variability. These parameters were then used in a discriminant analysis to identify individuals with a history of concussion with a classification accuracy of 94% [27]. It is important to note that the concussion history cohort was mostly asymptomatic by the time of testing, and many experienced their concussive injury several months prior [14,18,19,27]. Lastly, reaction time can also be used to identify individuals with a history of concussion, but only in elite athletes [19]. Reaction time did not significantly contribute to detecting concussion history in non-elite athletes or those participating in club sports. This finding suggests that different return to activity protocols may be needed for non-elite and professional athletes. Ultimately, concussion can negatively impact timing and accuracy aspects of motor performance, especially when under cognitive load, across a variety of injury severities and time since injury.

Our unique approach extends kinematic performance analyses by using systems identification techniques to quantify the contributions of sensorimotor memories to reach performance (Eq 1). This technique reveals the presence and relative contributions of trial-by-trial adaptive behavior despite subjects anecdotally reporting they cannot adapt to an unpredictable environment [8,16,20,25,29]. Recently, we have shown that this technique is predominantly an implicit process rather than explicit. Presumably, unpredictable perturbations discourage the use of explicit strategies in this task. When asking subjects to self-assess their reach errors after every movement (explicit memory recall), subjects report errors that are biased closer to a goal target and with less variability compared to actual reach errors [16]. We therefore assumed that actual reach error would be a proxy for implicit memory since it was not accurately recalled. Fitting a model that includes both explicitly-recalled errors and actual (implicitly maintained) errors revealed that the contribution of the actual errors significantly outperformed contributions from recalled errors, suggesting that the brain was predominantly using information inaccessible to the individual (i.e., implicit memory) [16]. We further supported this finding by having participants intentionally sabotage performance during the experiment in an attempt to manipulate how sensorimotor memories were used in adaptation. Remarkably, despite markedly reduced kinematic performance, the relative trial-by-trial contributions of sensorimotor memories did not change [8], further supporting the notion that this technique is probing an implicit process of adaptation.

Both the present study and the dual-dissociation cognitive-motor integration task described above [14] present kinematic parameters averaged across many trials to quantify performance between non-concussed and concussed individuals. A key similarity between the present study’s task employing memory-based trial-by-trial adaptation and Sergio et al.’s cognitive-motor integration task [14] is the involvement of implicit resources. The cognitive-motor integration task utilizes implicit spatial realignment when dissociating the plane of physical movement and visual feedback [30]. This implicit recalibration occurs slowly and alleviates the cognitive load of a rule-based explicit strategy. The present study’s trial-by-trial task quantifies how memories (some of which may be implicit) are used to reduce error on subsequent attempts (i.e., adaptation). It has recently been shown that averaged across-trial timing and accuracy measures can be intentionally sabotaged by emulating concussion symptoms despite the absence of injury in the past year [8]. Sabotaging performance may be exploited by athletes during pre-season baseline testing such that sabotaged scores appear comparable to post-concussion scores – thus impeding a concussion diagnosis and removal from play to avoid further injury. Sergio and colleagues have addressed this issue by instrumenting participants with a commercial four-channel EEG headset and using a neural network to detect sabotaged performance [31] which can then be used to justify a retest. We have addressed this issue through an inherent property of the implicit sensorimotor resources we are probing – implicit sensorimotor memories do not seem to be accessible or modifiable by the individual [8,16]. Methods to influence the contributions of sensorimotor memories during unpredictable perturbations include manipulations to the environment, such as providing or withholding visual feedback during reaches [16,20].

The present study has provided initial evidence that recent concussion can result in changes in trial-by-trial adaptation, but how sensorimotor memories contribute to that adaptation remains unclear. Sensorimotor memories are used to generate predictions of the upcoming perturbation, and these signals have been shown to correlate with activity in several cortical areas as well as in output pathways of the cerebellum and basal ganglia [32]. A recent study using multimodal MRI found significant correlations between concussion symptom recovery and changes in cerebellum grey matter susceptibility (thought to be from inflammation after impact), diffusivity of the corpus callosum, and connectivity of the caudate with the pallidum and thalamus [33] – regions thought to contribute to the adaptive control of actions [34]. Consequently, it is surprising that we did not see prominent effects of concussion on the relative contributions of motor memories to sensorimotor adaptation. However, strong conclusions are precluded by limiting factors of the present study which must be addressed before more definitive conclusions can be asserted.

### Limitations and Possibilities

A contemporary issue of concussion assessment stems from an assumption of honest, best-effort, baseline performance. Athletes have been known to underreport concussion symptoms [10] and may intentionally sabotage their baseline performance to levels typically seen after concussion – thus avoiding a concussion diagnosis. However, simple kinematic end-point measures such as reaction time, movement time, and performance variability can be intentionally manipulated when instructed or motivated to do so [8]. The robotic assessment of sensorimotor adaptation to unpredictable perturbations has demonstrated that it is inherently resistant to sabotaged performance [8] since this form of adaptation is predominantly implicit and discourages the use of explicit information. Participants can still sabotage kinematic performance, but they cannot sabotage how they use memories to produce sabotaged performance, thus providing an objective and unbiased measure of adaptive behavior.

The limited effects of concussion on reach performance and the modeling of sensorimotor adaptation may be driven by two major limitations identified within the present study: the heterogeneity of the concussed cohort and an observed practice effect within the robotic assessment of sensorimotor adaptation. Our concussed cohort of subjects was not only small, but also diverse with regards to the number of days between the injury and the first experimental visit. The average number of days before the first visit was about 6 days (see Fig 2A). The median time athletes spend in recovery before they become asymptomatic is typically 6 days [4]. We suspected that some subjects may have already significantly recovered by the time of their first visit which would reduce any effect size induced by subjects who had not yet recovered. Indeed, this suspicion was confirmed after splitting the concussed group by CogState’s composite assessment of reaction time. However, a major consequence of subdividing our concussed cohort was the resulting very small sample size of five concussed subjects. Our concussed cohort was composed of university students including non-athletes, intramural athletes, and some collegiate athletes which may have also contributed to the heterogeneity in our results.

The second major limitation was the presence of a practice effect that was evident in the control group for both kinematic measures and memory contribution coefficients. Any effects seen in the concussed group would likely contain the confounding practice effect in addition to the effect from the injury. Computer simulations based on model coefficients derived from healthy subjects suggest that during the first visit, subjects attempted to reduce steady-state error while allowing greater errors to arise from transients (changes) in the perturbations (Fig 8). However, by the second visit, subjects appeared to transition into reducing transient error at the expense of increased steady-state error. Put another way, subjects appeared to prioritize accuracy over precision on the first session, but by the second session, they improved precision at the expense of accuracy.

### Future Directions

The issue of the heterogenous concussed cohort can be mitigated by imposing tighter inclusion criteria and a more streamlined recruitment protocol. This includes recruiting only athletes within a set level of competition (i.e., collegiate athletes or intramural) which can be accomplished through partnerships with local athletic departments. These are also the individuals that will benefit the most from the robotic assessment for detecting concussion. This framework would also be beneficial in mitigating the latency between injury and the first testing visit. Several non-athlete subjects anecdotally reported that they had attempted to rest for a few days after sustaining their injury before seeking clinical care, where they were then invited to participate in the study. Several subjects continued to delay between solicitation and contacting the research team. Athletic trainers can significantly contribute to rectifying these issues if they are the point of contact for soliciting participants. Athletic trainers are typically present during the concussive injury and assess the athlete. As a member of the healthcare team as well as a member of the research team, the athletic trainer could potentially recruit the individual, thus removing steps the concussed individual must take to participate in research.

The second notable limitation was the presence of a practice effect during the robotic assessment of sensorimotor adaptation, which confounds the potential effect from concussion. The simplest way to mitigate this effect would be to include two baseline assessments prior to any potential post-concussion testing. The first baseline test would be discarded to account for confounding learning effects. The second test would act as the healthy pre-injury baseline. If a concussion were suspected, a third test could be administered; the significance of differences in performance variables such as reaction time and reach error, as well as differences in memory model coefficients could be assessed using a multilinear regression model with interaction terms [8].

Additional sensitivity to the effects of concussion on performance might also be attained by increasing task difficulty [27]. by. In one example, Brown, Sergio, and colleagues observed significantly longer movement times in concussed individuals when they had to perform a more difficult task, wherein target capture movements were performed in the horizontal plane but visual feedback was presented in the vertical plane such that hand movements to the right resulted in cursor movements to the left (i.e., two levels of visuomotor dissociation) [27].

Although the assessment of sensorimotor adaptation in the present study did not provide visual feedback, providing visual feedback during reaching changes the way individuals use sensorimotor memories to reduce reach errors on subsequent reaches [8,16]. Future research should explore the extent to which the robotic assessment of sensorimotor memories may or may not become more sensitive to concussion when additional neural resources and pathways supporting the visual control of movement are recruited.

The robotic assessment described above is unique to the field of concussion assessment in that it provides an objective and unbiased assessment of how memories and sensorimotor information are used to reduce subsequent reach error. These resources are predominantly implicit and are robust against attempts to sabotage performance. While concussion does impact motor performance and adaptation, it is unclear whether concussion significantly impacts how memories are used during adaptation. Future work will need to focus on a larger and/or more homogenous concussed cohort before generalizing to all sources and severity of concussion.

## Acknowledgements

We thank the participants in this study and the Marquette University Medical Clinic for assisting in subject recruitment.

## References

1. Sarmiento K, Thomas KE, Daugherty J, Waltzman D, Haarbauer-Krupa JK, Peterson AB, et al. Emergency Department Visits for Sports- and Recreation-Related Traumatic Brain Injuries Among Children — United States, 2010–2016. MMWR Morb Mortal Wkly Rep. 2019;68: 237–242. doi:10.15585/mmwr.mm6810a2

2. McCrory P, Meeuwisse W, Dvořák J, Aubry M, Bailes J, Broglio S, et al. Consensus statement on concussion in sport—the 5th international conference on concussion in sport held in Berlin, October 2016. Br J Sports Med. 2018;51: 838–847. doi:10.1136/bjsports-2017-097699

3. Goldberg LD, Dimeff RJ. Sideline management of sport-related concussions. Sports Med Arthrosc Rev. 2006;14: 199–205. doi:10.1097/01.jsa.0000212326.23560.09

4. McCrea M, Broglio S, McAllister T, Zhou W, Zhao S, Katz B, et al. Return to play and risk of repeat concussion in collegiate football players: comparative analysis from the NCAA Concussion Study (1999-2001) and CARE Consortium (2014-2017). Br J Sports Med. 2020;54: 102–109. doi:10.1136/bjsports-2019-100579

5. Boden BP, Tacchetti RL, Cantu RC, Knowles SB, Mueller FO. Catastrophic head injuries in high school and college football players. Am J Sports Med. 2007;35: 1075–81. doi:10.1177/0363546507299239

6. Elbin RJ, Sufrinko A, Schatz P, French J, Henry L, Burkhart S, et al. Removal From Play After Concussion and Recovery Time. Pediatrics. 2016;138: e20160910–e20160910. doi:10.1542/peds.2016-0910

7. Charek DB, Elbin RJ, Sufrinko A, Schatz P, D’Amico NR, Collins MW, et al. Preliminary Evidence of a Dose-Response for Continuing to Play on Recovery Time After Concussion. J Head Trauma Rehabil. 2020;35: 85–91. doi:10.1097/HTR.0000000000000476

8. Lantagne DD, Mrotek LA, Hoelzle JB, Thomas DG, Scheidt RA. Contribution of implicit memory to adaptation of movement extent during reaching against unpredictable spring-like loads: insensitivity to intentional suppression of kinematic performance. Exp Brain Res. 2023;241: 2209–2227. doi:10.1007/s00221-023-06664-z

9. McCrea M, Hammeke T, Olsen G, Leo P, Guskiewicz K. Unreported concussion in high school football players: implications for prevention. Clinical Journal of Sport Medicine. 2004;14: 13–17. Available: http://www.scopus.com/inward/record.url?eid=2-s2.0-25444503269&partnerID=40&md5=b0b9c245ac667d179068424985338b19

10. Meier TB, Brummel BJ, Singh R, Nerio CJ, Polanski DW, Bellgowan PSF. The underreporting of self-reported symptoms following sports-related concussion. J Sci Med Sport. 2015;18: 507–511. doi:10.1016/j.jsams.2014.07.008

11. Schatz P, Glatts C. “Sandbagging” Baseline Test Performance on ImPACT, Without Detection, Is More Difficult than It Appears. Archives of Clinical Neuropsychology. 2013;28: 236–244. doi:10.1093/arclin/act009

12. Higgins KL, Denney RL, Maerlender A. Sandbagging on the immediate post-concussion assessment and cognitive testing (impact) in a high school athlete population. Archives of Clinical Neuropsychology. 2017;32: 259–266. doi:10.1093/arclin/acw108

13. Raab CA, Peak AS, Knoderer C. Half of Purposeful Baseline Sandbaggers Undetected by ImPACT’s Embedded Invalidity Indicators. Arch Clin Neuropsychol. 2020;35: 283–290. doi:10.1093/arclin/acz001

14. Sergio LE, Gorbet DJ, Adams MS, Dobney DM. The Effects of Mild Traumatic Brain Injury on Cognitive-Motor Integration for Skilled Performance. Front Neurol. 2020;11: 1–9. doi:10.3389/fneur.2020.541630

15. Little CE, Dukelow SP, Schneider KJ, Emery CA. Using a Prism Paradigm to Identify Sensorimotor Impairment in Youth Following Concussion. J Head Trauma Rehabil. 2022;37: 189–198. doi:10.1097/HTR.0000000000000690

16. Lantagne DD, Mrotek LA, Slick R, Beardsley SA, Thomas DG, Scheidt RA. Contributions of implicit and explicit memories to sensorimotor adaptation of movement extent during goal-directed reaching. Exp Brain Res. 2021;239: 2445–2459. doi:10.1007/s00221-021-06134-4

17. Belanger HG, Vanderploeg RD. The neuropsychological impact of sports-related concussion: a meta-analysis. J Int Neuropsychol Soc. 2005;11: 345–57. Available: http://www.ncbi.nlm.nih.gov/pubmed/16209414

18. Dalecki M, Albines D, Macpherson A, Sergio LE. Prolonged congnitive-motor impairments in children and adolescents with a history of concussion. Concussion. 2016;1.

19. Hurtubise J, Gorbet D, Hamandi Y, Macpherson A, Sergio L. The effect of concussion history on cognitive-motor integration in elite hockey players. Concussion. 2016;1: CNC17. doi:10.2217/cnc-2016-0006

20. Judkins T, Scheidt RA. Visuo-proprioceptive interactions during adaptation of the human reach. J Neurophysiol. 2014;111: 868–887. doi:10.1152/jn.00314.2012

21. Scheidt RA, Reinkensmeyer DJ, Conditt MA, Zev Rymer W, Mussa-Ivaldi FA. Persistence of motor adaptation during constrained, multi-joint, arm movements. J Neurophysiol. 2000;84: 853–862. doi:10.1152/jn.2000.84.2.853

22. Maruff P, Thomas E, Cysique L, Brew B, Collie A, Snyder P, et al. Validity of the CogState brief battery: relationship to standardized tests and sensitivity to cognitive impairment in mild traumatic brain injury, schizophrenia, and AIDS dementia complex. Arch Clin Neuropsychol. 2009;24: 165–78. doi:10.1093/arclin/acp010

23. Cromer JA, Harel BT, Yu K, Valadka JS, Brunwin JW, Crawford CD, et al. Comparison of Cognitive Performance on the Cogstate Brief Battery When Taken In-Clinic, In-Group, and Unsupervised. Clinical Neuropsychologist. 2015;29: 542–558. doi:10.1080/13854046.2015.1054437

24. Louey AG, Cromer JA, Schembri AJ, Darby DG, Maruff P, Makdissi M, et al. Detecting cognitive impairment after concussion: Sensitivity of change from baseline and normative data methods using the CogSport/Axon cognitive test battery. Archives of Clinical Neuropsychology. 2014;29: 432–441. doi:10.1093/arclin/acu020

25. Scheidt RA, Dingwell JB, Mussa-Ivaldi FA. Learning to move amid uncertainty. J Neurophysiol. 2001;86: 971–85. doi:10.1152/jn.2001.86.2.971

26. van Donkelaar P, Osternig L, Chou L-S. Attentional and biomechanical deficits interact after mild traumatic brain injury. Exerc Sport Sci Rev. 2006;34: 77–82. doi:10.1249/00003677-200604000-00007

27. Brown JA, Dalecki M, Hughes C, Macpherson AK, Sergio LE. Cognitive-motor integration deficits in young adult athletes following concussion. BMC Sports Sci Med Rehabil. 2015;7: 1–12. doi:10.1186/s13102-015-0019-4

28. Dalecki M, Usand J, Van Gemmert AWA, Sergio LE. Motor Deficits in Youth with Concussion History: Issues with Task Novelty or Task Demand? Int J Sports Med. 2020;41: 688–695. doi:10.1055/a-1144-3217

29. Scheidt RA, Stoeckmann T. Reach Adaptation and Final Position Control Amid Environmental Uncertainty After Stroke. J Neurophysiol. 2007;97: 2824–2836. doi:10.1152/jn.00870.2006

30. Granek JA, Sergio LE. Evidence for distinct brain networks in the control of rule-based motor behavior. J Neurophysiol. 2015;114: 1298–1309. doi:10.1152/jn.00233.2014

31. Chaudhary M, Adams MS, Mukhopadhyay S, Litoiu M, Sergio LE. Sabotage Detection Using DL Models on EEG Data From a Cognitive-Motor Integration Task. Front Hum Neurosci. 2021;15: 662875. doi:10.3389/fnhum.2021.662875

32. Scheidt RA, Zimbelman JL, Salowitz NMG, Suminski AJ, Mosier KM, Houk J, et al. Remembering forward : Neural correlates of memory and prediction in human motor adaptation. Neuroimage. 2012;59: 582–600. doi:10.1016/j.neuroimage.2011.07.072

33. Pinky NN, Debert CT, Dukelow SP, Benson BW, Harris AD, Yeates KO, et al. Multimodal magnetic resonance imaging of youth sport-related concussion reveals acute changes in the cerebellum, basal ganglia, and corpus callosum that resolve with recovery. Front Hum Neurosci. 2022;16: 976013. doi:10.3389/fnhum.2022.976013

34. Houk JC, Davis JL, Beiser DG. Models of Information Processing in the Basal Ganglia. Cambridge, MA, USA: The MIT Press.; 1995.

